# Large scale gene duplication affected the European eel (*Anguilla anguilla*) after the 3R teleost duplication

**DOI:** 10.1101/232918

**Authors:** Christoffer Rozenfeld, Jose Blanca, Victor Gallego, Víctor García-Carpintero, Juan Germán Herranz-Jusdado, Luz Pérez, Juan F. Asturiano, Joaquín Cañizares, David S. Peñaranda

**Affiliations:** Grupo de Acuicultura y Biodiversidad. Instituto de Ciencia y Tecnología Animal. Universitat Politècnica de València. Camino de Vera s/n, 46022 Valencia, Spain; Instituto de Conservación y Mejora de la Agrodiversidad Valenciana, Universitat Politècnica de València, Camino de Vera 14, 46022, Valencia, Spain.

**Keywords:** European eel, PHYLDOG, 4dTv, whole genome duplication

## Abstract

Genomic scale duplication of genes generates raw genetic material, which may facilitate new adaptations for the organism. Previous studies on eels have reported specific gene duplications, however a species-specific large-scale gene duplication has never before been proposed. In this study, we have assembled a *de novo* European eel transcriptome and the data show more than a thousand gene duplications that happened, according to a 4dTv analysis, after the teleost specific 3R whole genome duplication (WGD). The European eel has a complex and peculiar life cycle, which involves extensive migration, drastic habitat changes and metamorphoses, all of which could have been facilitated by the genes derived from this large-scale gene duplication.

Of the paralogs created, those with a lower genetic distance are mostly found in tandem repeats, indicating that they are young segmental duplications. The older eel paralogs showed a different pattern, with more extensive synteny suggesting that a Whole Genome Duplication (WGD) event may have happened in the eel lineage. Furthermore, an enrichment analysis of eel specific paralogs further revealed GO-terms typically enriched after a WGD. Thus, this study, to the best of our knowledge, is the first to present evidence indicating an Anguillidae family specific large-scale gene duplication, which may include a 4R WGD.

## Introduction

Large-scale gene duplications can originate from one single event, like a whole genome duplication (WGD; Ohno, 1970) or from multiple smaller segmental duplication events (SDs; Gu et al. 2002). Any of these duplication events may contribute to species radiation, since both provide raw material for new genetic variation (Cañestro et al. 2013; Gu et al. 2002; Ohno 1970). It has been suggested that early in the vertebrate lineage two WGDs (1R and 2R) happened, resulting in species radiation and evolution of new traits (Cañestro et al. 2013; Dehal and Boore 2005; Gu et al. 2002; Ohno 1970). In teleosts, there is strong evidences to support an additional WGD, called the 3^rd^ teleost specific WGD (3R), which occurred in the base of the teleost lineage, between 350 and 320 million years ago (MYA; Aparicio et al. 2002; Christoffels et al. 2004; Howe et al. 2013; Vandepoele et al. 2004; Jaillon et al. 2004; Kasahara et al. 2007; Meyer and Peer 2005; Schartl et al. 2013). Previous studies have proposed that this extra 3R WGD is one of the possible causes of the massive species radiation observed in teleosts (Hoegg et al. 2004; Santini et al. 2009). In addition to 3R, multiple genus or species specific WGDs have been documented in teleosts, e.g. in salmonids (order Salmoniformes; Allendorf and Thorgaard, 1984; Johnson et al. 1987), sturgeons (order Acipenseriform; Ludwig et al. 2001), common carp (*Cyprinus carpio;* Larhammar and Risinger, 1994), goldfish (*Carassius auratus;* Ohno, 1970), suckers (family Catastomidae; Uyeno and Smith, 1972), and loaches (*Botia macracantha* and *Botia modesta;* Ferris and Whitt 1977).

As mentioned previously, other mechanisms to WGDs can create large-scale gene duplications. Several species have shown a high occurrence of relatively recent segmental duplications (SD), often found in tandem, with segments spanning from a few hundred base pairs to several genes e.g. in yeast (Llorente et al. 2000), daphnia (Colbourne et al. 2011), humans (Bailey et al. 2002; Gu et al. 2002; Vallente Samonte and Eichler 2016) and teleosts (Blomme et al. 2006; David et al. 2003; Jaillon et al. 2004; Lu et al. 2012; Rondeau et al. 2014). It is quite common for one of the copies of these SDs to get lost over time, possibly due to genetic drift or purifying selection. As a consequence, the genetic distance between two copies often tends to be quite small (Ohno, 1970). This process is known as the continuous mode hypothesis (Gu et al. 2002). In some cases however, these SDs have been conserved in high frequency at particular times, e.g. in yeast (Llorente et al. 2000), common carp (David et al. 2003) and humans (Asrar et al. 2013; Bailey et al. 2002; Gu et al. 2002; Hafeez et al. 2016). Some mechanisms, which could be facilitating this conservation include the processes of subfunctionalization, neofunctionalization or dosage selection (for review see Zhang, 2003). Furthermore, these processes have also been associated with the adaptation to new environments (Colbourne et al. 2011; Tautz and Domazet-lošo 2011).

The elopomorpha cohort, is one of the most basal teleost groups (Greenwood et al. 1966; Inoue et al. 2004). Elopomorphas are believed to originally be a marine species however, the 19 species of Anguillidae family, broke away from the ancestral trait and adapted a catadromous life style, migrating from their feeding grounds in freshwater rivers and lakes to their marine spawning grounds (Inoue et al. 2010; Munk et al. 2010; Schmidt 1923; Tsukamoto Katsumi, Nakai Izumi 1998). It is likely that species of the Anguillidae family originally performed relatively short reproductive migrations however, due to continental drift (Inoue et al. 2010; Tsukamoto et al. 2002) or changes in oceanic currents these migrations have since become vastly extensive (Jacobsen et al. 2014), with a total migrating distance of >6.000 km in the case of the European eel (Righton et al. 2016).

Several previous studies have revealed a high occurrence of duplicated genes in eels (Dufour et al. 2005; Henkel et al. 2012; Lafont et al. 2016; Maugars and Dufour 2015; Morini et al. 2015; Pasqualini et al. 2009; Pasquier et al. 2012; Rozenfeld et al. 2016; Morini et al. 2017). E.g. Lafont et al. (2016) found two paralog genes of *ift140, tleo2, nme4, xpo6,* and *unkl,* in the *gper* genomic regions of the eel. Only one copy of these genes has been observed in other teleosts. These results led Lafont et al. (2016) to hypothesize i) that the whole region containing *gper* could have been duplicated in *Anguilla* eels, and maybe also in other teleosts, and ii) that the retention of duplicated genes may be higher in eels than in other teleosts.

For the present study, we assembled a *de novo* European eel transcriptome from Illumina RNA sequencing data. In order to study species-specific duplications and the timings of the events that created them, we ran phylogenetic reconstructions and calculated fourfold synonymous third-codon transversion (4dTv) distances. This analysis was performed on our transcriptome, and on multiple other fish transcriptomes and genomes. Our analysis revealed a high accumulation of duplicated genes in eel compared to other teleost species (which do not have a confirmed 4R duplication event in their linage). Many of these duplications are restricted to the eel lineage, and the 4dTv analyses suggested that these duplications happened much later than the 3R WGD shared by all teleosts. We will discuss in more depth if these eel specific duplications are the result of several SDs or one eel-specific 4R WGD. To our knowledge, this is the first published evidence of a large-scale lineage specific duplication in the elopomorpha cohort.

## Results

### Transcriptome assemblies and genomes

To assemble a *de novo* European eel transcriptome, we performed high quality RNA extractions from forebrain, pituitary, and testis samples, of one eel, following the protocol described by Peña-Llopis and Brugarolas (2013). The RNA was then quality tested on the Bio-Rad Bioanalyser, which yielded average RIN values of 8.90. From this RNA, in total 181 million Illumina reads, with a length of 101 bp, were produced. These reads were assembled by using the Trinity assembler after a digital normalization step that left 75 million representative reads. The transcriptomes of Northern pike (*Esox lucius*), elephantnose fish (*Gnathonemus petersii*) and silver arowana (*Osteoglossum bicirrhosum*) were also assembled by Trinity using Illumina reads from the Phylofish database (Pasquier et al. 2016). The resulting unigenes were clustered by using a transitive clustering approach to create sets of very similar transcripts. The number of unigenes (henceforth referred to as transcripts) assembled ranged from 68489 to 78610 (table 1) and the number of transcript clusters from 49154 to 55667 (henceforth referred to as genes; table 2).

**Table 1.** Size and quality of included transcriptomes from: European eel (*Anguilla Anguilla*), Northern Pike (*Esox Lucius*), elephantnose fish (*Gnathonemus petersi*), and silver arowana (*Osteoglossum bicirrhosum).*

| Species | N.º Reads | Q30 | Transcripts |
| --- | --- | --- | --- |
| European eel | 181322106 | 0.994 | 77247 |
| Northern Pike | 553710218 | 0.989 | 68489 |
| Elephantnose fish | 498451616 | 0.993 | 74642 |
| Silver arowana | 490649254 | 0.992 | 78610 |

**Table 2.** Quantities of included genes per included species: European eel (*Anguilla anguilla*), Zebrafish (*Danio rerio*), Northern pike (*Esox Lucius*), Elephantnose fish (*Gnathonemus petersi*), Spotted gar (*Lepisosteus oculatus*), Silver arowana (*Osteoglossum bicirrhosum*), Atlantic salmon (*Salmo salar*), Fugu (*Takifugu rubripes*), and Platyfish (*Xiphophorus maculatus).*

| Species | Transcripts | Genes | Representative transcripts | Representative transcripts with predicted protein | Gene family transcripts | % of genes assigned to a gene family |
| --- | --- | --- | --- | --- | --- | --- |
| European eel | 77247 | 54879 | 54845 | 27696 | 25862 | 93.38 |
| Zebrafish | 58274 | 32189 | 32189 | 25790 | 22703 | 88.03 |
| Northern pike | 68489 | 49154 | 49154 | 23843 | 21696 | 90.99 |
| Elephantnose fish | 74642 | 50455 | 50455 | 24857 | 22036 | 88.65 |
| Spotted gar | 22483 | 18341 | 18341 | 18341 | 17872 | 97.44 |
| Silver arowana | 78610 | 55667 | 55667 | 24938 | 21604 | 86.63 |
| Atlantic salmon | 109584 | 55104 | 55104 | 48593 | 42625 | 87.72 |
| Fugu | 47841 | 18523 | 18523 | 18523 | 17698 | 95.55 |
| Platyfish | 20454 | 20379 | 20379 | 20379 | 19807 | 97.19 |

The genomes of zebrafish (*Danio rerio*), northern pike, spotted gar (*Lepisosteus oculatus*), fugu (*Takifugu rubripes*), and platyfish (*Xiphophorus maculatus*) were obtained from the ENSEMBL database, the Atlantic salmon ( *Salmo salar*) genome was downloaded from NCBI and the published eel genome was downloaded from the ZF-genomics web site (Henkel et al. 2012).

### Genome and transcriptome quality assessment

In order to test the completeness of the transcriptomes and genomes we ran a BUSCO analysis in which we looked for a set of single-copy orthologues, typically found in fish genomes (Simão et al. 2015; Fig. 1). In general, genomes were more complete than transcriptomes according to the BUSCO assessment, and were thus preferred. However, in the cases of the pike and eel, the transcriptomes outperformed the genomes (Fig. 1), and these transcriptomes were therefore used for further analysis.

**Figure 1.**
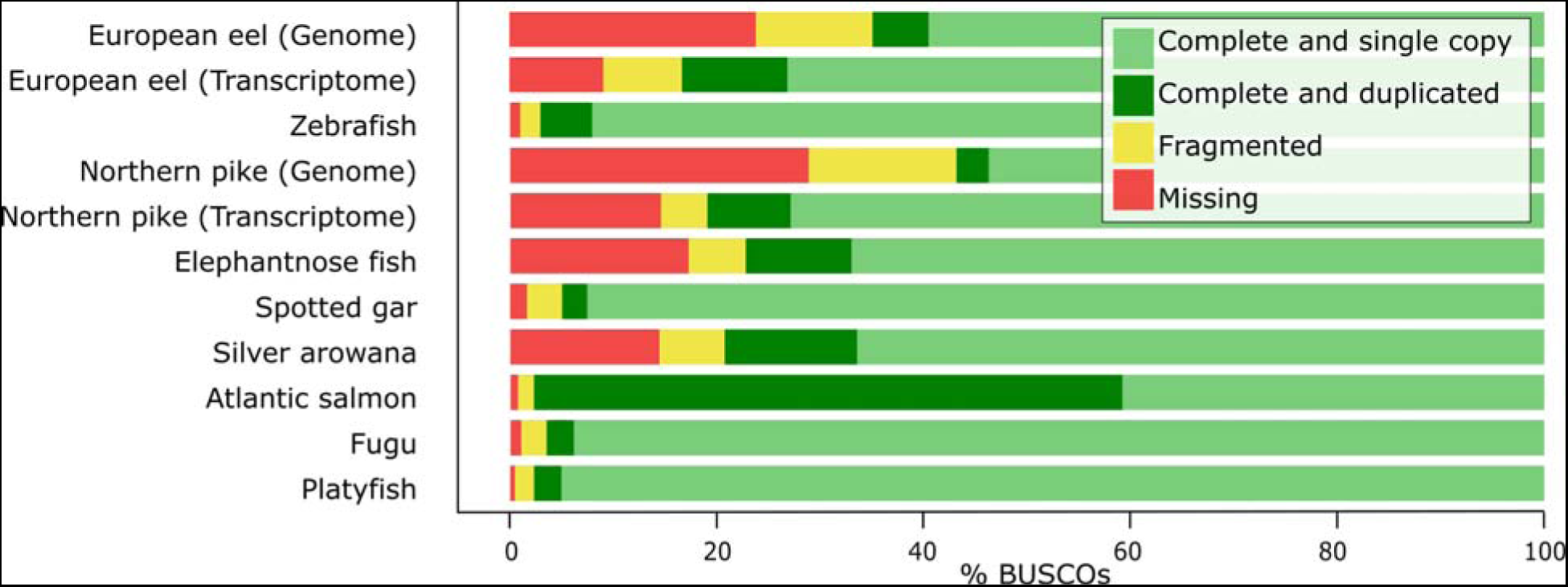
BUSCO (Benchmarking set of Universal Single-Copy Orthologues) result for every genome and transcriptome, one per row. The sequence of a BUSCO gene can be found complete or fragmented in each genome and it can be found once (single copy), more than once (duplicated) or not found (missing). Included genomes: European eel (*Anguilla anguilla*), zebrafish (*Danio rerio*), northern pike (*Esox lucius*), spotted gar (*Lepisosteus oculatus*), fugu (*Takifugu rubripes*), platyfish (*Xiphophorus maculatus*) and Atlantic salmon (*Salmo salar).* Included transcriptomes: European eel, northern pike, elephantnose fish (*Gnathonemus petersii*) and silver arowana (*Osteoglossum bicirrhosum).*

In order to further test the completeness of the eel and pike transcriptomes, we mapped the eel and pike RNA-seq reads to the transcriptome assembly using BWA-MEM (Li and Durbin 2010) and to the genome using the software HISAT2 (Pertea et al. 2016). The percentages of reads that mapped concordantly against the genome and the transcriptome were 65.8 and 91.9% respectively for eel, and 44.6 and 85.8% for pike. Likewise, previous published eel RNA-sequencing experiments were also mapped to the eel genome and transcriptome. In this case, 52.2% (Coppe et al. 2010), 57.9% (Burgerhout et al. 2016), and 66.18% (Ager-Wick et al. 2013) reads mapped concordantly against the eel genome whereas 84.3% (Coppe et al. 2010), 69.5% (Burgerhout et al. 2016), and 87.32 % (Ager-Wick et al. 2013) mapped against the transcriptome.

### Gene families

One representative transcript for each gene and species was selected; the longest one for genomes and the most expressed one for transcriptomes. The OrthoMCL web service (Li et al. 2003) assigned gene families to the genes. The percentage of genes assigned to a family ranged from 88.0% (zebrafish) to 97.4% (spotted gar; table 2). Overall, 17003 gene families were covered, from which 13823 protein and codon alignments were built. These families contained between 2 and 161 genes, with 9 genes per family being the mode (suppl. Fig. 1).

### Phylogenetic reconstruction and duplication dating

PHYLDOG (Boussau et al. 2013) was run 10 independent times using 8,000 protein alignments chosen at random. Overall, PHYLDOG created trees for 10,352 gene families and, based on the tree topology, it labelled the branches in which gene duplication events had happened. All 10 runs produced a species tree that matched the species tree topology created by phylobayes (Lartillot et al. 2009) with a CAT-GTR model built with the concatenation of 100 protein alignments and a Neighbour-Joining tree built with a 4dTv distance matrix (Fig. 2).

**Figure 2.**
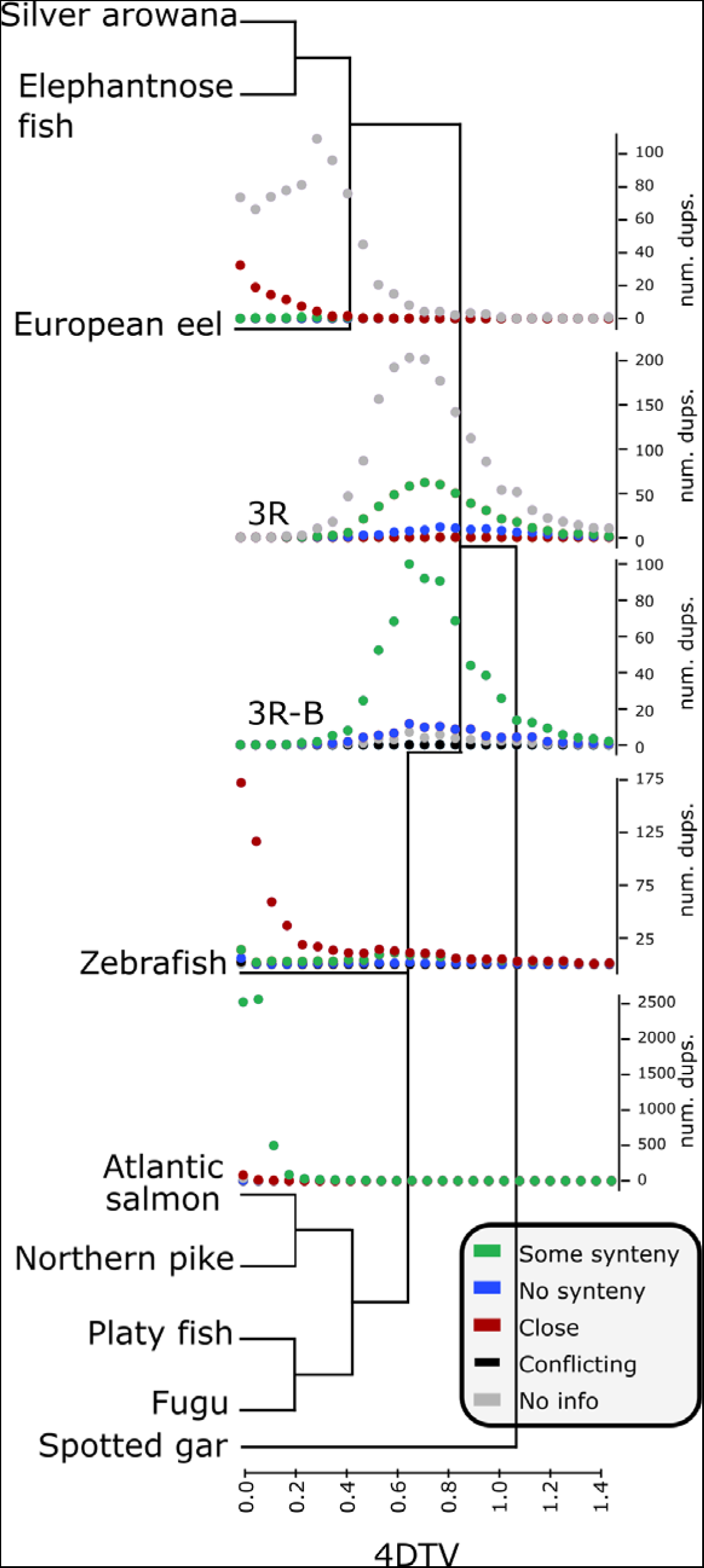
Species cladogram generated by PHYLDOG for the species included in this study: European eel (*Anguilla anguilla*), zebrafish (*Danio rerio*), northern pike (*Esox lucius*), spotted gar (*Lepisosteus oculatus*), fugu (*Takifugu rubripes*), platyfish (*Xiphophorus maculatus*), Atlantic salmon (*Salmo salar*), elephantnose fish (*Gnathonemus petersii*) and silver arowana (*Osteoglossum bicirrhosum*). PHYLDOG also determined the duplication events and for each of these events the 4dTv and the synteny type found around the gene was determined. Only the 4dTv distributions for the branches with most duplications are represented over the corresponding cladogram branch, for the distributions for all branches refer to supplementary figure 2. The synteny types are the following: close, the copies originated by the duplication are close in the genome; some synteny, some genes close to the one duplicated are also found to be duplicated close by; no synteny, there are no paralogs for others genes found close to the paralog copies created by the duplication; no information, the duplicated genes are located in small scaffolds with not enough genes close by; conflicting syntenies, different synteny classification found in the genomes of the different species affected by the duplication)

For each duplication found in each gene family tree, the 4dTv distance between the genes was calculated, and by grouping them according to the species tree branch in which it happened, the distribution of the 4dTvs for each lineage was built. Each 4dTv distribution was further divided according to the synteny type found in the region where the paralogs of each gene family were located (Fig. 2 and suppl. Fig. 2). The duplications were thus labeled according to the genomic region where the resulting paralogs were found. In some cases the paralog pairs were found close to each other (labelled as close), denoting a tandem SD, in other cases they were in syntenic regions where paralogs from other gene families were also located (labelled as "some synteny”), possibly denoting a WGD and, finally, in some other cases, there were not enough close genes (labelled as "no info”) in the genome assembly or conflicting evidence was found (labelled as “conflicting syntenies”).

PHYLDOG assigned 4,308 duplications to the basal teleost branch, after the split of the spotted gar (Fig. 2 and Fig. 3), with a 4dTv mode of 0.8. Of the paralogs created by these duplications 63.1% were located in regions with some synteny, 2.4% were close to each other, and 32.3% had no synteny. These percentages are calculated without taking into account the duplications where no information regarding the physical location of the genes could be established. The following branch (directly following the split of the eel, arowana and elephantnose fish) was assigned 1,525 duplications and showed very similar distributions with an overall 4dTv mode of 0.75. The eel specific branch was assigned 1460 duplications of which 16.5, 75.8, and 7.2% were labelled as some synteny, close and without synteny, respectively. Notably, most of the eel specific duplications lacked sufficient physical genomic location information. The duplications that generated close genes in tandem within the eel genome clearly showed a different distribution to the ones located in syntenic regions and the ones with no information. The tandem ones tended to be more recent according to their 4dTv (fig. 2) while the syntenic ones, and most of the duplications without sufficient genomic location information, showed a 4dTv mode of 0.4. In both the salmon and zebrafish specific branches most duplications seemed quite recent, according to their 4dTv values, with 8,712 and 1,452 branch specific duplications, respectively. In the case of the salmon, most duplications (80.6%) were characterized by paralogs located in syntenic regions whereas most zebrafish paralogues (54.6%) were close tandem SDs (Fig. 2 and Fig. 3).

**Figure 3.**
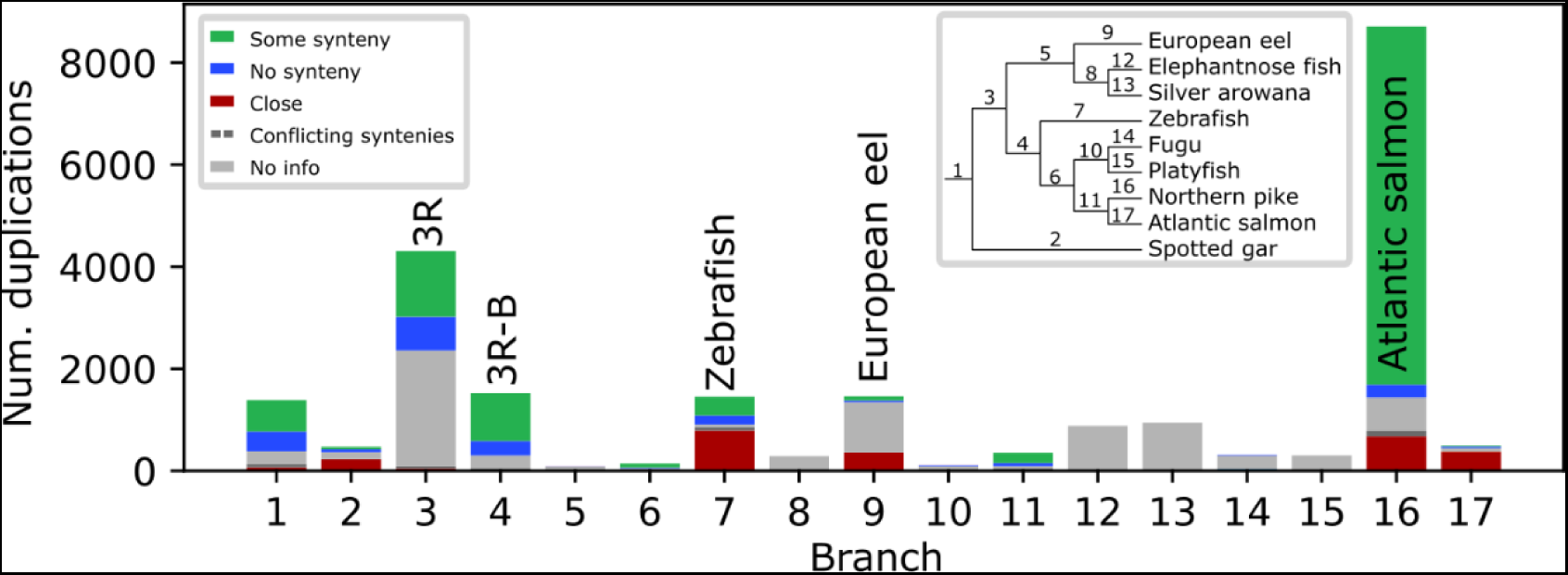
Barplot of the number of duplications PHYLDOG assigned to each branch of the species tree. Bars are numbered according to the cladogram in the upper righthand corner. 3R indicates the branch where the 3R teleost-specific whole genome duplication is hypothesized to have happened. 3R-B indicates the basal branch of the remaining teleosts after the split of the elopomorphas and osteoglossomorphas. Each bar is subdivided into the synteny types described in figure 2.

In order to investigate the timing of the main eel duplication event in greater depth, we compared the 4dTv distribution found for eel paralogs with the 4dTv distribution built for the eel orthologs with elephantnose fish, and arowana. The results showed a 4dTv maximum of 0.4 for the main eel paralog peak, and 0.5 for the peaks corresponding to the speciation event that separated elephantnose fish and arowana from eel (Fig. 4).

**Figure 4.**
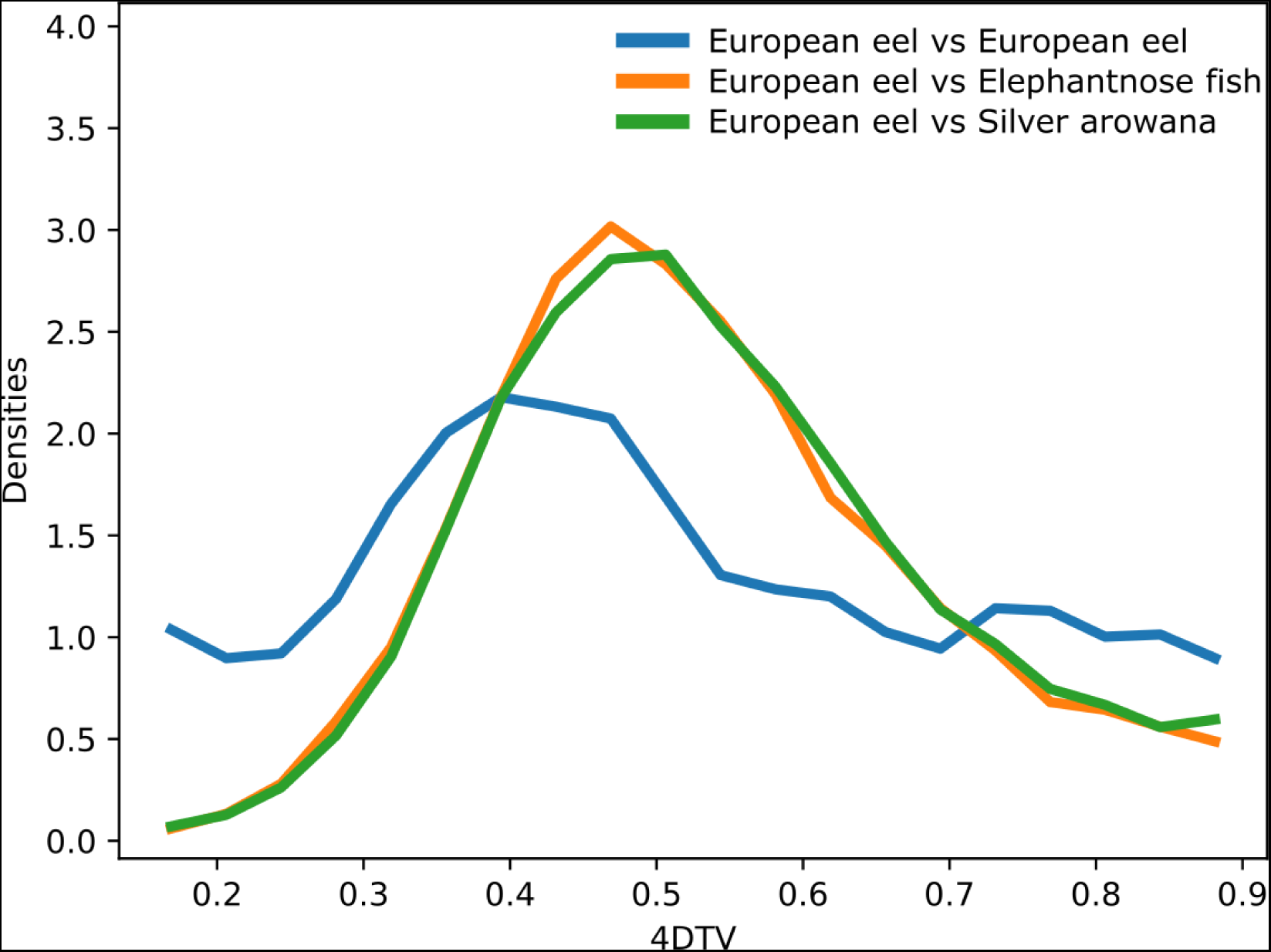
4dTv distribution of European eel (*Anguilla anguilla*) paralogs (blue), European eel and elephantnose fish (*Gnathonemus petersii*) orthologs (green), and European eel and silver arowana (*Osteoglossum bicirrhosum*) orthologs (orange). This analysis was carried out in the gene families without taking into account the PHYLDOG result.

### Investigation of functional category enrichment

To investigate if some functional categories were overrepresented in the eel duplications, an enrichment test was carried out. GO terms were assigned to 9570 gene families by comparing them to the annotated EggNOG gene families (Huerta-Cepas et al. 2016). The GO annotation can be found in suppl. table 1. From these terms we performed an enrichment analysis with the topGO R library (Alexa and Rahnenfuhrer 2016).

The resulting enriched GO-terms, are presented in the suppl. table 2. In most cases these terms are involved either in signal transduction (GTPase, MAPK, sphingosine-1-phosphate), morphological alterations (convergent extension involved in axis elongation and gastrulation, heart development, and pronephric glomerulus development, pigmentation), or forebrain development.

Additionally, KEGG terms were assigned to 16466 eel genes using BlastKOALA (Kanehisa et al. 2016) and the terms related to the genes involved in the eel duplication were mapped onto the KEGG pathways using the KEGG Mapper tool. A fisher test, corrected for multiple testing using False Discovery Rate, was used to look for enriched KEGG pathways in the eel branch (Suppl. Table 3). Most of the KEGG pathways found to be enriched were related to immune system, nervous system, oocyte, apoptosis, cell adhesion, amino acid metabolism, glycan biosynthesis, and signal transduction. Several key genes related to the immune system were also duplicated, including: Cytohesin-associated scaffolding protein (*casp*) c-jun N-terminal knase (*jnk*) and vav proto-oncogene (*vav*), involved in the lymphocyte differentiation and activation, the interleukine and T-cell receptors (*tcr’s*), major histone compatibility complex (*mch*) I, *mch* II, and several cytokines including *tgf*-beta (transforming growth factor beta) and B-cell activating factor (*baff*). There were also numerous duplications related to apoptosis, including activator protein 1 (*ap-1*), c-fos proto-oncogene (*c-fos*), *casp*, serine protease (*omi*), mitochondrial Rho GTPase (*miro*), Death executioner (*bcl-2*), Caspase Dredd, X-linked inhibitor of apoptosis protein (*xiap*), *casp8*, and Baculoviral inhibitor-of-apoptosis repeat (*bir*). In the nervous system tyrosine hydroxylase (*th*) and monoamine oxidase, as well as some receptors (*gaba-a, gaba-b, ampa, mglur, crfr1,* and *girk*) were duplicated, as well as several genes involved in synaptic exocytosis (*rab3a, munc-18,* syntaxin, *vgat*), and Golf and Transductin, which are involved in olfaction and phototransduction. In the oocyte more affected genes were found, including the progesterone receptor (*pgr*), early mitotic inhibitor 2 (*emi2*), aurora-B phosphatase (*glc7*), cullin F-box containing complex (*scf-skp*), insulin-like growth factor 1 (*igf1*), and cytoplasmic polyadenylation element binding protein (*cpeb*).

## Discussion

The observed data has shown, for the first time to our knowledge, that in the eel lineage more than a thousand gene families have genes that are the result of a large-scale gene duplication that happened after the teleost specific 3R WGD. Only Atlantic salmon, due to its salmonid-specific 4R WGD, showed a higher amount of conserved duplicates, whereas the number of duplicated gene families found in eel and zebrafish were similar. Furthermore, the paralog 4dTv distribution shows a unique pattern in the case of the eel branch when compared to the other teleosts.

The duplications assigned by the phylogenetic analysis to the eel specific branch, after the split of arowana and elephantnose fish, showed a distribution of 4dTv distances younger than those corresponding to the 3R duplication and older than the salmonid duplications. This result was replicated in an independent analysis, not based on phylogenetic tree topologies. In this case the distributions of the 4dTv distances (Fig. 4) were calculated between the eel genes found in the gene families (eel paralogs) and between the eel sequences and the arowana and elephantnose fish sequences (orthologs). Both the phylogenetic topologies and the 4dTv distances corroborate the hypothesis that the main eel duplication happened after the teleost specific 3R duplication event (320-350 MYA; Vandepoele et al. 2004 and Christoffels et al. 2004), and after the arowana and elephantnose fish split, but before the salmonid duplication (80-100 MYA; Macqueen et al. 2014). Given the 4dTv distribution found, we could assume that this duplication event is shared by all members of the Anguilla genus, as they are estimated to first appear 20-50 MYA (Minegishi et al. 2005). To be more precise about the timing of the main eel duplication event it would be advisable to study other elopomorpha transcriptomes or genomes to analyze if they share the same duplication.

Not all the PHYLDOG duplication assignments to the basal branches of the species tree are as trustworthy as the lineage specific ones. According to PHYLDOG the duplications usually associated with the 3R WGD would be split into two events, one would correspond to the 3R branch and the other with the one that gave rise to zebrafish, salmon and fugu (indicated as 3R-B; Fig. 2). If PHYLDOG is right and there were really two genome duplications the 4dTv distributions of the events should be different. These distributions are quite similar, although not completely identical. Nevertheless, duplications this old are almost completely saturated in terms of 4dTv, decreasing the accuracy of the measurement. Alternatively, the PHYLDOG split of the 3R duplication could be explained as an artifact. Only 3 species make up the daughter clade of the 3R branch at one side of the split and some gene families will only include representatives from the species of the other daughter clade. This could make the phylogenetic based assignment of the gene duplication to a particular branch more error prone. Thus, some gene family duplications created by 3R might end up wrongly assigned by PHYLDOG to 3R-B. This might have happened even with some zebrafish lineage specific duplications. The 4dTv distribution for the zebrafish branch shows a bump that overlaps with the 3R duplication. That might be due to a real old duplication in the zebrafish lineage, but it is also possible that it is a PHYLDOG artifact. The zebrafish genome is among the most complete genomes, with the most well-supported protein coding gene annotation, used in this study. Therefore, it might be the case that some duplicated genes could be found only in this species making the PHYLDOG assignment more difficult.

In previous studies, data consistent with an eel specific duplication event has been reported, although it has never been interpreted in this way. Recently, the transcriptome of several teleosts was sequenced (Pasquier et al. 2016), and the European eel was the species with the highest number of contigs, expect for species with a documented 4R WGD in their lineage. In the additional data included by Inoue et al. (2015), in their analysis of the gene loss process that followed the teleost 3R WGD, both the eel and zebrafish are the species with the highest percentage of duplications (36.6 and 31.9%, respectively). Thus, these two studies are in concordance with our analysis.

Previously, several lineage specific gene duplications have been found and studied in eel, in studies focusing on particular genes. For instance, Morini et al. (2015) found that the leptin receptors were duplicated in the eel. However, their phylogeny did not include other basal teleosts and was compatible with both the hypothesis that those genes were duplicated in an eel specific duplication event or in the teleost 3R WGD. Similarly, Lafont et al. (2016) found several species-specific duplicated genes in the eel genome, but they attributed it to less extensive loss of genes in the eel lineage after the 3R duplication. Furthermore, several other analyses of single genes have reached the same conclusion (Dufour et al. 2005; Maugars and Dufour 2015; Pasqualini et al. 2009; Pasquier et al. 2012; Morini et al. 2017; Henkel et al. 2012). In general, these studies were based on tree topologies that did not include other basal teleost species and often did not report the genetic distances.

The eel duplication was also not detected in the analysis of the published eel genome (Henkel et al. 2012). This genome was quite fragmented and, according to our BUSCO analysis, it was more incomplete than our transcriptome, thus including less duplications (Fig. 1). Moreover, no global 4dTv or Ks distribution was calculated, but a remarkably high number of Hox genes (73) were found, thus these results are also compatible with our current analysis.

In the proposed species phylogeny (Fig. 2) the eel is located in a basal position as found in other previous phylogenies. The location of the osteoglossomorphes order, represented by the elephantnose fish and arowana is, however, in disagreement with previous phylogenies based on elopomorpha mitochondrial genomes (Inoue et al. 2003) and on the nuclear arowana genome (Austin et al. 2015). In these phylogenies arowana is the basal node to the teleosts, whereas it is grouped with the eel in ours. In the transcriptome based phylogeny proposed by the PhyloFish project (Pasquier et al. 2016) the topology is reversed and the Anguilliformes appear as the basal teleosts whereas the Osteoglossiformes appear to have split more recently.

The branch that separated eel and arowana, in the arowana genome paper, was one of the shortest and had a lower posterior probability than the others; this could be the reason why these different phylogenies disagree. This disparity in the results is unlikely to be due to the phylogenetic methods used. In this study we have used the neighbor joining, maximum likelihood and bayesian approaches, and all of them agree that arowana and eel form two sister clades. The difference might be due to the extra species that we are including, the elephantnose fish. To be more confident of the topology of these events in the base of the teleosts, it would be advisable to include other basal species.

Two different mechanisms can create genomic scale duplications: WGDs or many small-scale SDs, a process referred to as the continuous mode hypothesis (Gu et al. 2002). The latter process has been observed in many species, including yeast (Llorente et al. 2000), fruit flies (Zhou et al. 2008), water fleas (Colbourne et al. 2011), humans (Bailey et al. 2002; Gu et al. 2002; Vallente Samonte and Eichler 2016), several plant species (Cui et al. 2006) and teleosts (Blomme et al. 2006; David et al. 2003; Jaillon et al. 2004; Lu et al. 2012; Rondeau et al. 2014). Usually, most of these SDs are lost soon after they are generated. Therefore, in the genome, many young paralog pairs, and few old, can be found. In a 4dTv distribution this pattern would be detected as a peak of 4dTv values with a mode close to 0. Moreover, SDs are usually the result of tandem duplications, therefore very similar paralogs located in tandem are likely to be the result of this process. Patterns compatible with this scenario were seen in several species in our analysis, including: zebrafish, elephantnose fish, arowana, pike, fugu, spotted gar, salmon and eel (suppl. Fig. 2). The high amount of SDs that we detected in zebrafish has been previously documented (Blomme et al. 2006; Lu et al. 2012). In some other species it has been shown that these SDs could be retained at specific points in time, possibly during specific evolutionary events e.g. in yeast (Llorente et al. 2000), common carp (David et al. 2003) and humans (Bailey et al. 2002; Hughes et al. 2001). These events have been linked to the adaptation of a species to a new environment (Chain et al. 2014; Colbourne et al. 2011; Tautz and Domazet-lošo 2011).

In eel, the 4dTv distribution pattern found is quite distinct as it reflects two modes; one of younger and one of older duplications. Of these duplications, the older ones are clearly older than those found, for example, in zebrafish or salmon. Furthermore, the genomic surroundings of the paralogs of the younger duplications are quite different from those of the older duplications. The younger duplications, which have a lower 4dTv, tend to be located close together in the genome and are likely to have been generated recently by tandem SDs, whereas the older duplications are not usually found in tandem. A WGD should have left behind blocks of syntenic regions similar to those detected in our analysis for the 3R teleost and the salmon WGD. In the case of the eel, an increase in these syntenic blocks was also detected in the older duplications. However most genomic regions are very fragmented in the genome assembly and thus, we lack physical genomic location information for many genes. However, from the evidence available we can hypothesize that most of the older duplications are likely to be the result of a WGD which occurred in the eel lineage, and that an analysis with the latest eel genome assembly published (Jansen et al. 2017), but not available without restrictions, would detect more syntenic regions. In other words, the numerous duplications found in eel are likely to have been generated by a WGD followed by many SDs, a pattern which has also been observed in primates (Gu et al. 2002), and common carp (David et al. 2003).

In this study, several gene function analyses were carried out to study overrepresented functions among the eel specific duplications. These overrepresentations could be linked to several adaptations that have taken place throughout eel evolution, e.g. the inclusion of a leptocephali larvae stage to their life history (Inoue et al. 2004), the adaptation to a catadromous lifecycle (Inoue et al. 2010), and the adaptation to withhold maturation until after the extensive reproductive migration (Righton et al. 2016; van Ginneken et al. 2005). Other mechanisms which perhaps influence the conservation of paralogs are: dosage selection (Glasauer and Neuhauss 2014) and segregation avoidance (Hahn 2009). It has been suggested that these mechanisms conserve duplicated genes related to specific biological processes, such as development, signaling, ion transport, metabolism and neuronal function after WGDS (Berthelot et al. 2014; Blomme et al. 2006; Brunet et al. 2006; Kassahn et al. 2009).

Specifically, 54 GO-terms were found to be enriched among the eel specific duplications (suppl. table 2). Interestingly, several of the GO-terms found to be enriched among the eel specific duplications form part of some of the aforementioned processes, including; development, ion transport, signaling, neuronal function, and metabolism. The high number of enriched GO-terms, which are part of processes that are often conserved after WGDs, suggests that the duplication event here described is a WGD. It is likely that they have been conserved due to the mechanisms regulating gene conservation after WGD rather than due to specific necessities of the *Anguilla* species. Other GO terms that were found to be duplicated in eel are not usually found in other WGDs, for example “pigmentation”. As the eel has incorporated several pigmentation alterations into its lifecycle, it is likely that the genes associated with this GO-term are conserved due to new functions acquired in the *Anguilla* species. Most of the pigmentation changes undergone by the eel are linked to the transition between the marine and freshwater environment, therefore the duplication of these genes might have generated the necessary raw genetic material for adaptation to the catadromous lifecycle.

Furthermore, 54 KEGG pathways were also found to be enriched among the eel specific duplications (suppl. table 3). As in the case of the GO-terms, several of these pathways are involved in signaling, metabolism and neuronal function. Additionally, olfactory transduction and several pathways involved in immune response e.g. Tuberculosis, Th1 and Th2 cell differentiation, Bacterial invasion of epithelial cells, Th17 cell differentiation, and others were found to be enriched. Lu et al. (2012) also found that immune response pathways and olfactory receptor activity were enriched among the recent segmental duplications found in zebrafish. Several studies have also found immune response genes to be enriched among other recent SDs (Conrad and Antonarakis 2007; Kasahara et al. 2007; She et al. 2008; Stein et al. 2007; Wang et al. 2012). Thus this enrichment in immune response genes could be linked to eel SDs. However, in these studies, the recent duplications were found to be mostly in the components interacting with pathogens, possibly to contribute to the response against different pathogens, as opposed to the components downstream of the receptors, which make up most of the enriched pathways found in our study. Also, among the eel specific duplications were the progestin receptors which have recently been characterized in eel (Morini et al. 2017). Progestins are known as maturation-inducing steroids promoting sperm maturation and spermiation (for review see Scott et al. 2010), and the two paralogs do show differential expression during maturation (Morini et al. 2017).

The most significantly enriched pathway found in the eel duplications is the dopaminergic synapse pathway. Dopamine (DA) is an essential neurotransmitter in vertebrates, with several functions (Davila et al. 2003; Hsia et al. 1999). DA has been proven to be important in teleosts, where DA has been found to have an inhibitory role on the gonadotropic activity of the pituitaries (Dufour et al. 2005; Peter et al. 1986). In the case of the eel in particular, DA inhibition completely arrests puberty before their oceanic migration (Vidal et al. 2004), indicating that DA has a much stronger inhibitor effect in eel compared to most other teleosts (Dufour 1988; Vidal et al. 2004). This suggests that the duplicated genes involved in the dopaminergic synapse pathway may have been conserved during the adaptation to block maturation until after the extensive reproductive migration. Among the duplicated genes assigned to the dopaminergic synapse pathway, we found tyrosine hydroxylase (TH). TH is the rate limiting enzyme of the DA biosynthesis (Nagatsu et al. 1964), and is therefore often used as an indicator of DA tone in eel (Davila et al. 2003; Weltzien et al. 2015). As genes are rarely conserved without a specific function or necessity, the presence of two TH genes in the eel encourages suspicion of potential differential expression or function between the two, which may prove important for the regulation of the DA induced inhibition of puberty observed in pre-migration eels.

In conclusion, the data presented strongly suggest that a vast amount of genes have been duplicated specifically in the eel lineage. Furthermore, the synteny, 4dTv, and enrichment analyses suggest that these genes derive both from a WGD as well as continuously created SDs, and that they are related to the eel specific physiology. To our knowledge this is thus the first evidence published suggesting a possible eel lineage specific 4R WGD.

## Materials and methods

### Fish husbandry

Ten immature farm eel males (mean body weight 96.7±3.6 g±SEM) supplied by Valenciana de Acuicultura S.A. (Puzol, Valencia, Spain) were transported to the Aquaculture Laboratory at the Universitat Politècnica de València, Spain. The fish were kept in a 200-L tank, equipped with individual recirculation systems, a temperature control system (with heaters and, coolers), and aeration. The fish were gradually acclimatized to sea water (final salinity 37 ± 0.3%o), over the course of two weeks. The temperature, oxygen level and pH of rearing were 20 °C, 7-8 mg/L and ~ 8.2, respectively. The tank was covered to maintain, as much as possible, a constant dark photoperiod and the fish were starved throughout the holding period. After acclimation, the fish were sacrificed in order to collect samples of forebrain, pituitary, and testis tissues.

### Human and Animal Rights

This study was carried out in strict accordance with the recommendations given in the Guide for the Care and Use of Laboratory Animals of the Spanish Royal Decree 53/2013 regarding the protection of animals used for scientific purposes (BOE 2013), and in accordance with the European Union regulations concerning the protection of experimental animals (Dir 86/609/EEC). The protocol was approved by the Experimental Animal Ethics Committee from the Universitat Politècnica de València (UPV) and final permission was given by the local government (Generalitat Valenciana, Permit Number: 2014/VSC/PEA/00147). The fish were sacrificed using anesthesia and all efforts were made to minimize suffering.

### RNA extraction and sequencing

High quality RNA was extracted from forebrain, pituitary, and testis samples following the protocol developed by Peña-Llopis and Brugarolas (2013). Quantity and quality were tested on a Bio-Rad Bioanalyser (Bio-Rad Laboratories, Hercules, CA, USA), selecting the samples with RIN values and amounts higher than >8 >3 μg of total RNA, respectively. Total RNA samples were shipped to the company Macrogen Korea (Seoul, South Korea). Then, a mRNA purification was carried out using Sera-mag Magnetic Oligo (dT) Beads, followed by buffer fragmentation. Reverse transcription was followed by PCR amplification to prepare the samples for sequencing. The strand information was kept in an Illumina Hiseq-2000 sequencer (Illumina, San Diego, USA). Resulting raw sequences are available at the NCBI Sequence Read Archive (SRA) as stated in the section titled “Data accessibility”.

### Transcriptome assemblies and genomes

The software FastQC (Andrews 2010) was used to assess the quality of the raw reads generated by Macrogen. Thereafter, trimmomatic (Bolger et al. 2014) was used to trim the reads, eliminating known adaptor sequences, and low quality regions. Finally, trimmed reads shorter than 50 bp were filtered out. Eel reads were digitally normalized before assembly by khmer software (Crusoe et al. 2015) using a k-mer length of 25 and a coverage of 100. Further, The RNA-Seq raw reads for pike, arowana and elephantnose fish were downloaded from the PhyloFish project (Pasquier et al. 2016). All transcriptomes were then assembled using Trinity software (Haas et al. 2013), with the read orientation and sense (in the eel case) into account. The transcripts assembled were filtered according to their complexity (with a DUST score threshold of 7 and a DUST window of 64), length (with a minimum length of 500 bp), and level of expression (with a TPM threshold of 1). After assembly, the CDSs and proteins were annotated using the Trinotate functional annotation pipeline (Haas et al. 2013).

Transcripts that share k-mers are clustered by Trinity, however, these transcripts might correspond to different transcript forms of the same gene or to closely related genes from a gene family. We split these transcripts into genes by running a transitive clustering based on a blast search. In this clustering transcripts which shared at least 100 bp with a minimum identity of 97% were considered to be isoforms of the same gene. Thus, some Trinity clusters were split into several genes. For each gene, the most expressed transcript, according to Salmon (Patro et al. 2017), was chosen as its representative.

The available eel genome was downloaded from the ZF-Genomics web site (Henkel et al. 2012). The salmon genome assembled by the International Cooperation to Sequence the Atlantic Salmon Genome was downloaded from NCBI (Lien et al. 2016). The genomes of zebrafish (Howe et al. 2013), fugu (Kai et al. 2011), spotted gar (Braasch et al. 2016), and platyfish (Schartl et al. 2013) were downloaded from ENSEMBL (release 87). The pike genome (Rondeau et al. 2014) was downloaded from the Northern Pike Genome web site (Genbank accession GCA_000721915.1). For each gene in the genomes, the longest transcript was chosen as the representative.

### Genome and transcriptome quality assessment

In order to check the quality of the transcriptomes and genomes we looked for the BUSCO conserved gene set in them (Simão et al. 2015). BUSCOs are conserved proteins, and are expected to be found in any complete genome or transcriptome. Therefore, the number of present, missing, or fragmented BUSCOs can be used as a quality control of a genome or transcriptome assembly. For this assessment the Actinopterygii (*odb9*) gene set, which consists of 4584 single-copy genes that are present in at least 90% of Actinopterygii species was used. As an additional comparison between the transcriptome and genomes of pike and eel, the RNA-seq reads were mapped both to the genome and transcriptome assemblies using the softwares HISAT2 (Pertea et al. 2016) and BWA-MEM (Li and Durbin 2010), respectively.

### Gene families

Genes were clustered into gene families by the OrthoMCL web service (Li et al. 2003). For each gene family a multiple protein alignment was built. To avoid transcriptome assembly artifacts proteins longer than 1,500 amino acids, transcripts with a DUST score higher than 7 or sequences with more than 40% of gaps in the alignment were filtered out. The software Clustal Omega (Sievers et al. 2011) carried out the protein multiple alignment and trimAl (Capella-Gutiérrez et al. 2009) removed the regions with too many gaps or those difficult to align. The protein alignment was used as a template to build the codon alignment by aligning the transcript sequences against the corresponding protein using the protein2dna exonerate algorithm (Slater and Birney 2005).

### Phylogenetic reconstruction and duplication dating

The resulting protein alignments were used by PHYLDOG (Boussau et al. 2012) software to generate a species tree as well as a family tree corresponding to each alignment. Due to the high memory requirements of PHYLDOG not all gene families could be run in the same analysis so 10 analyses were carried out, choosing 8000 protein alignments at random for each. Once all runs were finished, we checked that the species tree topology of all the 10 species trees, matched exactly. . PHYLDOG uses a maximum likelihood approach to simultaneously coestimate the species and gene family trees from all individual alignments. From the topology of the gene family trees, it is capable of inferring when the duplications in each family happened. Alternatively, the species phylogeny was also reconstructed using a bayesian approach by using PhyloBayes MPI version 1.7 (Lartillot et al. 2009). From the gene families that had one gene for each species, 100 were chosen at random to create a concatenated alignment of 43566 aminoacids. The model used was CAT-GTR and three independent MCMC chains were run for 39872, 56328, and 39285 iterations.

Finally, a neighbor joining tree based on the fourfold synonymous third-codon transversion distances (4dTv) was also calculated (Tang et al. 2008). Between any pair of sequences the number of transversions found in the third base of the codon was divided by the number of fourfold degenerated codons. A correction to the 4dTv was applied: ln(1 - 2 * distance) / -2. For each pair of species a 4dTv distance was calculated. 4dTvs were calculated between the sequences of those species found in each gene family codon alignment. The distribution of those 4dTvs was fitted with a log normal mixture model using the scikit-learn Gaussian Mixture class. The number of components required was one for all the species pairs, except for those where a bimodal distribution was found due to a recent speciation event. The distance between any two species was the mode of the fitted model. The neighbor joining tree was built using the BioPython Tree Construction class. This process is implemented in the fit_fdtv_distributions module found in the Python scripts (suppl. Material 1).

The 4dTv was calculated for each duplication tagged by PHYLDOG within any gene family. A duplication event defines a subtree in the gene family tree, and this subtree defines two child branches, so the 4dTv calculated for that event was the mean of the 4dTv between all combinations of sequences found between those branches. These calculations are implemented by the functions calculate_4dTv and calculate_mean_fdtv_for_tree found in the Python scripts (suppl. Material 1).

### Synteny

Furthermore, the kind of event that created each duplication was characterized by analyzing the conserved synteny between the paralogs created by that duplication within a particular genome. A duplication may derive from a SD that could have occurred in tandem or not, or from a WGD, among others. Tandem SDs would create paralogs found close to each other in the genome, whereas the paralogs created by a WGD would be far away, but surrounded by similar genes in each of the duplicated regions. We also have to consider that several phylogenetically close species can be affected by the same older duplication event. Therefore, these traces of the duplication event could be found in the genomes of those different species and, if no other genomic rearrangement happened since, these traces should match each other and convey the same information. With this in mind, we categorized duplications as one of 4 classes: i) the paralog genes that were found close to each other in the genome, within a 50 gene distance were labelled as close, ii) the paralogs which were found in syntenic regions where 2 or more paralogous from other gene families were located within a 50 genes distance, not necessarily in the same colineal order, were labelled as "some synteny”, iii) the cases in which fewer than 2 gene families could be identified within a 50 gene distance from both of the paralogous genes were labelled as "no info”, and iv) the cases in which conflicting evidence was found in the genomes of the different species affected by the duplication were labelled as “conflicting syntenies”. This labelling of the duplications was carried out by the Python function determine_if_pair_is_close_or_synthenic and the Python class GenomeLocator, found in the scripts (suppl. material 1). The location of each gene in a genome was obtained by performing a BLAST search with its representative transcript against the genome.

### Investigation of functional category enrichment

The EggNOG database has GO annotations for each of its gene families (Huerta-Cepas et al. 2016). To match our gene families with those from the EggNOG database, the protein sequence with least gaps for each of our families was selected and a HMMER search (Finn et al. 2011) was carried out against the EggNOG position weight matrices with an e-value threshold of 0.0001. The GO annotation of the best EggNOG hit in this search was transferred to our family. The enrichment analysis was carried out using the fisher statistic and the weight algorithm of the topGO library (Alexa and Rahnenfuhrer 2016) from the bioconductor project. The R script go_enrichment_analysis found in the scripts (suppl. Material 1) implements this analysis. Eel transcripts were annotated using the BlastKOALA KEGG service (Kanehisa et al. 2016) and a fisher exact test was carried out, using the scipy implementation, to look for overrepresented KEGG pathways in the eel duplications.

## Data accessibility

The raw RNA-sequencing reads from brain, pituitary, and testis samples from European eel (*Anguilla anguilla*) have been deposited at GenBank (http://www.ncbi.nlm.nih.gov/genbank) under accession no. XX. Deposition and acquiring of accession number was not finished at the time of first submission but will be settled within a few days.

## Acknowledgements

This study received funding from the project REPRO-TEMP (AGL2013-41646-R) funded by the Spanish Ministry of Economy and Competitiveness, and from the European Union’s Horizon 2020 research and innovation program under the Marie Skłodowska-Curie grant agreement No 642893 (IMPRESS). V. Gallego has a postdoc grant from the UPV (PAID-10-16).

